# A small-molecule TrkB ligand improves dendritic spine phenotypes and atypical behaviors in female Rett syndrome mice

**DOI:** 10.1101/2023.11.09.566435

**Authors:** Destynie Medeiros, Karen Ayala-Baylon, Hailey Egido-Betancourt, Eric Miller, Christopher Chapleau, Holly Robinson, Mary L. Phillips, Tao Yang, Frank M. Longo, Wei Li, Lucas Pozzo-Miller

## Abstract

Rett syndrome (RTT) is a neurodevelopmental disorder caused by mutations in methyl-CpG-binding protein-2 (*MECP2*), encoding a transcriptional regulator of many genes, including brain-derived neurotrophic factor (*Bdnf*). BDNF mRNA and protein levels are lower in RTT autopsy brains and in multiple brain regions of *Mecp2*-deficient mice, and experimentally increasing BDNF levels improve atypical phenotypes in *Mecp2* mutant mice. Due to the low blood-brain barrier permeability of BDNF itself, we tested the effects of a brain penetrant, small molecule ligand of its TrkB receptors. Applied *in vitro*, LM22A-4 increased dendritic spine density in pyramidal neurons in cultured hippocampal slices from postnatal day (P) 7 male *Mecp2* knockout (KO) mice as much as recombinant BDNF, and both effects were prevented by the TrkB receptor inhibitors K-252a and ANA-12. Consistent with its partial agonist activity, LM22A-4 did not affect spine density in CA1 pyramidal neurons in slice cultures from male wildtype (WT) mice, where typical BDNF levels outcompete its binding to TrkB. To identify neurons of known genotypes in the ‘mosaic’ brain of female *Mecp2* heterozygous (HET) mice, we treated 4–6-month-old female MeCP2-GFP WT and HET mice with peripheral injections of LM22A-4 for 4 weeks. Surprisingly, mutant neurons lacking MeCP2-GFP showed dendritic spine volumes comparable to that in WT controls, while MeCP2-GFP-expressing neurons showed larger spines, similar to the phenotype we described in symptomatic male *Mecp2* KO mice where all neurons lack MeCP2. Consistent with this non-cell-autonomous mechanism, a 4-week systemic treatment with LM22A-4 had an effect only in MeCP2-GFP-expressing neurons in female *Mecp2* HET mice, bringing dendritic spine volumes down to WT control levels, and without affecting spines of MeCP2-GFP-lacking neurons. At the behavioral level, we found that female *Mecp2* HET mice engaged in aggressive behaviors significantly more than WT controls, which were reduced to WT levels by a 4-week systemic treatment with LM22A-4. Altogether, these data revealed differences in dendritic spine size and altered behaviors in *Mecp2* HET mice, in addition to provide support to the potential usefulness of BDNF-related therapeutic approaches such as the partial TrkB agonist LM22A-4.

## INTRODUCTION

Rett syndrome (RTT) is an X chromosome-linked neurodevelopmental disorder with associated intellectual disabilities that affects approximately 1:10,000 females worldwide (Katz et al., 2012; Neul and Zoghbi, 2004). The majority of RTT individuals carry loss-of-function mutations in the gene that encodes methyl-CpG-binding protein-2 (MeCP2), a transcriptional modulator that binds to methylated DNA sites (Amir et al., 1999; Chahrour et al., 2008; Nan et al., 1997; Percy and Lane, 2005).

The transcription, synthesis, intracellular transport, and activity-dependent release of the neurotrophin brain-derived neurotrophic factor (BDNF) are impaired in a number of neurological disorders including RTT and Huntington’s disease (Gines et al., 2010; Hartmann et al., 2012; Tapia-Arancibia et al., 2008). The relevance of BDNF deficiency to RTT pathogenesis is supported by the observations that *Bdnf* expression is directly regulated by MeCP2 in an activity-dependent manner (Chen et al., 2003; Martinowich et al., 2003; Zhou et al., 2006), BDNF levels are lower in multiple brain areas of *Mecp2*-deficient mice (Abuhatzira et al., 2007; Li et al., 2012; Schmid et al., 2012; Wang et al., 2006), and increasing BDNF levels via genetic or pharmacological manipulations improve some of the deficits observed in *Mecp2*-deficient neurons and mice (Chang et al., 2006; Chapleau et al., 2009; Ogier et al., 2007). BDNF signaling via its tropomyosin related kinase B (TrkB) receptor plays a key role in neuronal and synaptic development and adult synaptic plasticity, and *in vitro* application of BDNF increases dendritic spine density through its activation (Alonso et al., 2004; Tyler and Pozzo-Miller, 2001). In addition, hippocampal pyramidal neurons of *Mecp2* KO mice have lower dendritic spine density (Chapleau et al., 2012), impaired BDNF-induced membrane currents and Ca^2+^ signals mediated by TRPC3 channels (Li et al., 2012), and reduced trafficking and activity-dependent release of BDNF-GFP (Xu et al., 2014). While these studies indicate that BDNF deficiency is a key component in RTT pathogenesis (Katz, 2014; Li and Pozzo-Miller, 2014), the therapeutic potential of BDNF is limited by its low blood-brain barrier permeability and short plasma half-life (Poduslo and Curran, 1996).

An alternative to BDNF itself is to use synthetic small molecules that target the TrkB receptor as ligands. An established preclinical candidate is LM22A-4, a ‘mimetic’ of the BDNF loop-2 domain that activates TrkB and its downstream signaling pathways (Massa et al., 2010). Indeed, LM22A-4 improved disease phenotypes in mouse models of Huntington’s disease (Simmons et al., 2013), Dravet’s disease (Gu et al., 2022), and chemotherapy-induced cognitive decline (Geraghty et al., 2019). In female *Mecp2* HET mice, a 2-month treatment with LM22A-4 improved breathing irregularities (Kron et al., 2012; Schmid et al., 2012), and reduced network hyperactivity in hippocampal slices and restored long-term potentiation of excitatory synaptic transmission, as well as improved object location memory (Li et al., 2017). Similarly, a second generation TrkB ligand based on LM22A-4, also improved breathing patterns and motor deficits in *Mecp2* HET mice (Adams et al., 2020). In the present study, we tested the effects of LM22A-4 on dendritic spine density and size in hippocampal pyramidal neurons of female *Mecp2*-deficient mice, as well as in a machine-learning unbiased screen of open field behaviors in female *Mecp2* HET mice interacting with novel and familiar mice.

## RESULTS AND DISCUSSION

### LM22A-4 increases dendritic spine density in pyramidal neurons of hippocampal slice cultures from male *Mecp2* KO mice via activation of TrkB receptors

We first confirmed that eYFP-expressing hippocampal CA1 pyramidal neurons in organotypic slice cultures from postnatal-day 7 (P7) male *Mecp2* KO mice have lower spine density than neurons in cultured slices from age-matched male WT mice (p=0.0032, unpaired t-test; n=17 *Mecp2* KO neurons / 10 slices vs. n=14 WT neurons / 9 slices; Supplemental Figure 1), as we reported previously (Chapleau et al., 2012). In addition, the volume of individual spines is larger in CA1 pyramidal neurons in slice cultures from *Mecp2* KO mice (p<0.0001, Kolmogorov-Smirnov [K-S] test; n=1,458 *Mecp2* KO spines / 17 neurons / 10 slices vs. n=2,139 WT spines / 14 neurons / 9 slices; Supplemental Figure 1C), as we described in dissociated cultures of hippocampal neurons from P1 male *Mecp2* KO mice (Xu et al., 2017) and in CA1 pyramidal neurons in *ex vivo* slices from symptomatic P45-65 male *Mecp2* KO mice (Li et al., 2016).

Consistent with multiple reports in different experimental preparations, including ours in hippocampal slice cultures from neonatal rats (Alonso et al., 2004; Chapleau et al., 2012; Chapleau et al., 2008; Tyler and Pozzo-Miller, 2001), BDNF (250ng/ mL, 48hs) increased spine density in CA1 pyramidal neurons in slice cultures from both male *Mecp2* KO mice (p=0.0026, one-way ANOVA with Bonferroni’s post-hoc test; n=17 control *Mecp2* KO neurons / 10 slices vs. n=13 BDNF *Mecp2* KO neurons / 5 slices) and WT littermates (p=0.0042, ANOVA-Bonferroni’s; n=14 control WT neurons / 9 slices vs. n=15 BDNF WT neurons /5 slices; Supplemental Figure 1D). Also consistent with prior reports, these effects of BDNF were reduced by the non-selective Trk receptor inhibitor K-252a (200nM) in WT slice cultures (p=0.0088, ANOVA-Bonferroni’s; n=13 BDNF+K-252a WT neurons / 6 slices vs. n=15 BDNF WT neurons / 5 slices; Supplemental Figure 1D). Unexpectedly, K-252a did not alter the BDNF effect in *Mecp2* KO neurons (p>0.9999, ANOVA-Bonferroni’s; n=15 BDNF+K-252a *Mecp2* KO neurons / 6 slices vs. n=13 BDNF *Mecp2* KO neurons / 5 slices) and in fact increased spine density by itself (p<0.0001, ANOVA-Bonferroni’s; n= K-252a *Mecp2* KO neurons / 4 slices vs. n=17 control *Mecp2* KO neurons / 10 slices; Supplemental Figure 1D), which may reflect an altered signaling balance between the low affinity p75 receptor and TrkB (e.g. Chapleau and Pozzo-Miller, 2012) in the absence of MeCP2.

Consistent with the activation of TrkB, LM22A-4 (500 nM, 48hs) increased spine density in pyramidal neurons in areas CA1 and CA3 of hippocampal slice cultures from male *Mecp2* KO mice (p<0.0001, ANOVA-Bonferroni’s; n=17 control *Mecp2* KO neurons / 10 slices vs. n=10 LM22A-4 *Mecp2* KO neurons / 7 slices; Supplemental Figure 1E). Consistent with its partial agonism at TrkB receptors (Massa et al., 2010), and the lower levels of BDNF in *Mecp2* mutant mice compared to WT mice (Katz, 2014; Li and Pozzo-Miller, 2014), LM22A-4 had no effect on spine density in pyramidal neurons from male WT mice (p>0.9999, ANOVA-Bonferroni’s; n=12 control WT neurons / 11 slices vs. n=13 LM22A-4 WT neurons / 9 slices). Similar to the effect of BDNF described above, K-252a and the selective TrkB inhibitor ANA-12 (Cazorla et al., 2011) both reduced the effect of LM22A-4 on spine density in *Mecp2* KO neurons (p=0.0011, ANOVA-Bonferroni’s; n=10 LM22A-4 + ANA-12 *Mecp2* KO neurons / 7 slices vs. n=10 LM22A-4 *Mecp2* KO neurons / 7 slices; Supplemental Figure 1E). Similar to the spinogenic effect of K-252a, ANA-12 also increased spine density by itself in *Mecp2* KO neurons (p<0.0001, ANOVA-Bonferroni’s; n=11 ANA-12 *Mecp2* KO neurons / 5 slices vs. n=10 LM22A-4 *Mecp2* KO neurons / 7 slices; Supplemental Figure 1E).

### LM22A-4 modulates spine volume only in MeCP2-expressing CA1 pyramidal neurons in female heterozygous MeCP2-GFP mice

To identify neurons of known genotypes in the ‘mosaic’ brain of female *Mecp2* HET mice (due to X-chromosome inactivation), we crossed female *Mecp2* HET mice with male mice expressing GFP-tagged MeCP2 (Schmid et al., 2012). For quantitative analyses of dendritic morphology, CA1 pyramidal neurons in *ex vivo* hippocampal slices from young adult mice were identified as MeCP2-expressing or MeCP2-lacking based on the nuclear presence of GFP and were filled with biocytin through a whole-cell patch pipette, followed by confocal microscopy of Alexa-488-tagged streptavidin (Figure 1A).

**Figure 1:**
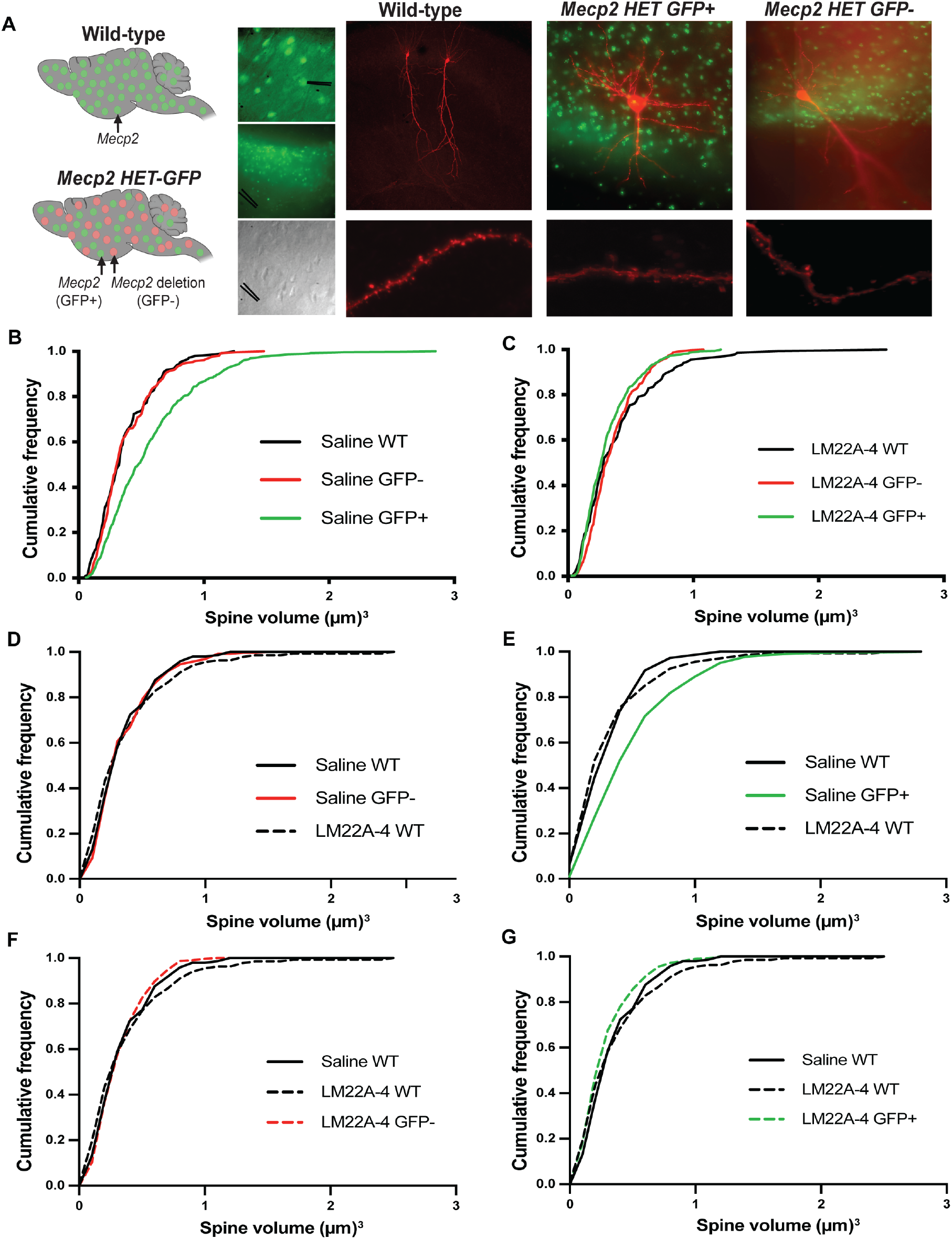
Systemic LM22A-4 reduces the volume of CA1 dendritic spines of MeCP2-expressing neurons in the ‘mosaic’ brain of female MeCP2-GFP HET mice, without affecting that of MeCP2-lacking neurons. **(A)** Left. Representative schematic of female WT brain with all neurons express GFP-tagged MeCP2 protein, and female MeCP2-GFP HET brain where *Mecp2*-expressing neurons express GFP-tagged MeCP2, and *Mecp2*-lacking neurons do not. Right, Representative images of biocytin filled CA1 pyramidal neurons in slices from female WT and female MeCP2-GFP HET mice. Bottom, Representative images of dendritic spines. **(B-G)** Cumulative probability distributions of individual spine volumes in CA1 pyramidal neurons in slices from WT and MeCP2-GFP HET mice treated with LM22A-4 or saline (control).

Consistent with the spine phenotype of symptomatic male *Mecp2* KO mice (P45-65), where their density is similar to that in age-matched male WT mice (Li et al., 2016), spine density is similar in the two cellular genotypes in 6 months-old female *MeCP2*-GFP HET mice, and comparable to that in age-matched female WT mice (p>0.9999, ANOVA-Bonferroni’s; n=35 MeCP2-expressing in HET neurons / 18 slices vs. n= 24 MeCP2-lacking in HET neurons / 14 slices vs. n=9 WT neurons / 6 slices; Supplemental Figure 2). Unexpectedly, MeCP2-expressing CA1 pyramidal neurons in female MeCP2-GFP HET mice have dendritic spines with larger volumes than neighboring MeCP2-lacking neurons and CA1 pyramidal neurons in slices from age-matched female *Mecp2* HET mice (p<0.0001, K-S test; n=521 spines / 35 neurons / 18 slices MeCP2-expressing in HET vs. n= 257 spines / 24 neurons / 14 slices MeCP2-lacking in HET vs. n=145 spines / 9 neurons / 6 slices WT; Figure 1B). This observation suggests that dendritic spines of MeCP2-expressing neurons in the ‘mosaic’ brain of female MeCP2-GFP HET mice retain their ‘sensitivity’ to neuronal activity (e.g. Segal et al., 2000) under the conditions of heightened hippocampal activity observed in female *Mecp2* HET mice (Li et al., 2017), which is comparable to that in the hippocampus of male *Mecp2* KO mice (Calfa et al., 2011; Calfa et al., 2015; Li et al., 2016).

Four-to-six-month-old female MeCP2-GFP HET mice and their age-matched WT littermates were treated with either LM22A-4 (50 mg/kg in saline, twice daily) or vehicle (saline) for 4 weeks, as described (Kron et al., 2012; Li et al., 2017; Schmid et al., 2012). Consistent with the phenotype described above, LM22A-4 did not affect spine density (p>0.9999, ANOVA-Bonferroni’s; n=48 MeCP2-expressing in LM22A-4 HET neurons / 24 slices vs. n=28 MeCP2-lacking in LM22A-4 HET neurons / 18 slices vs. p=0.0423, K-S test; n=11 LM22A-4 WT neurons / 9 slices; Supplemental Figure 2). However, LM22A-4 reduced spine volume only in MeCP2-expressing pyramidal neurons of female MeCP2-GFP HET mice, reaching values comparable to those in MeCP2-lacking neurons in control HET mice (p>0.9999, K-S test; n=505 spines / 48 neurons / 24 slices MeCP2-expressing in LM22A-4 HET vs. n= 257 spines / 24 neurons / 14 slices MeCP2-lacking in control HET) as well as those in female control WT mice (p>0.9999, K-S test; n=505 spines / 48 neurons / 24 slices MeCP2-expressing in LM22A-4 HET vs. n=145 spines / 9 neurons / 6 slices control WT; Figure 1C-G).

Similar to its actions *in vitro*, LM22A-4 had no effect on spine volume of female WT mice (p>0.9999, K-S test; n=134 spines / 11 neurons / 9 slices LM22A-4 WT vs. n=145 spines / 9 neurons / 6 slices control WT). Spine volume was similar in control WT mice, LM22A-4-treated WT mice, and MeCP2-lacking neurons in LM22A-4-treated HET mice (p>0.9999, K-S test; n= 145 spines / 9 neurons / 6 slices control WT vs. n= 134 spines / 11 neurons / 9 slices LM22A-4 WT vs. n=303 spines / 28 neurons / 18 slices LM22A-4 MeCP2-lacking; Figure 1F). Spine volumes of MeCP2-expressing neurons in LM22A-4-treated HET mice were partially rescued to levels comparable to those of control WT and LM22A-4 WT (p<0.0001, K-S test; n=505 spines / 48 neurons / 24 slices LM22A-4 MeCP2-expressing vs. n=134 spines / 11 neurons / 9 slices LM22A-4 WT vs. n=145 spines / 9 neurons / 6 slices control WT; Figure 2G).

**Figure 2:**
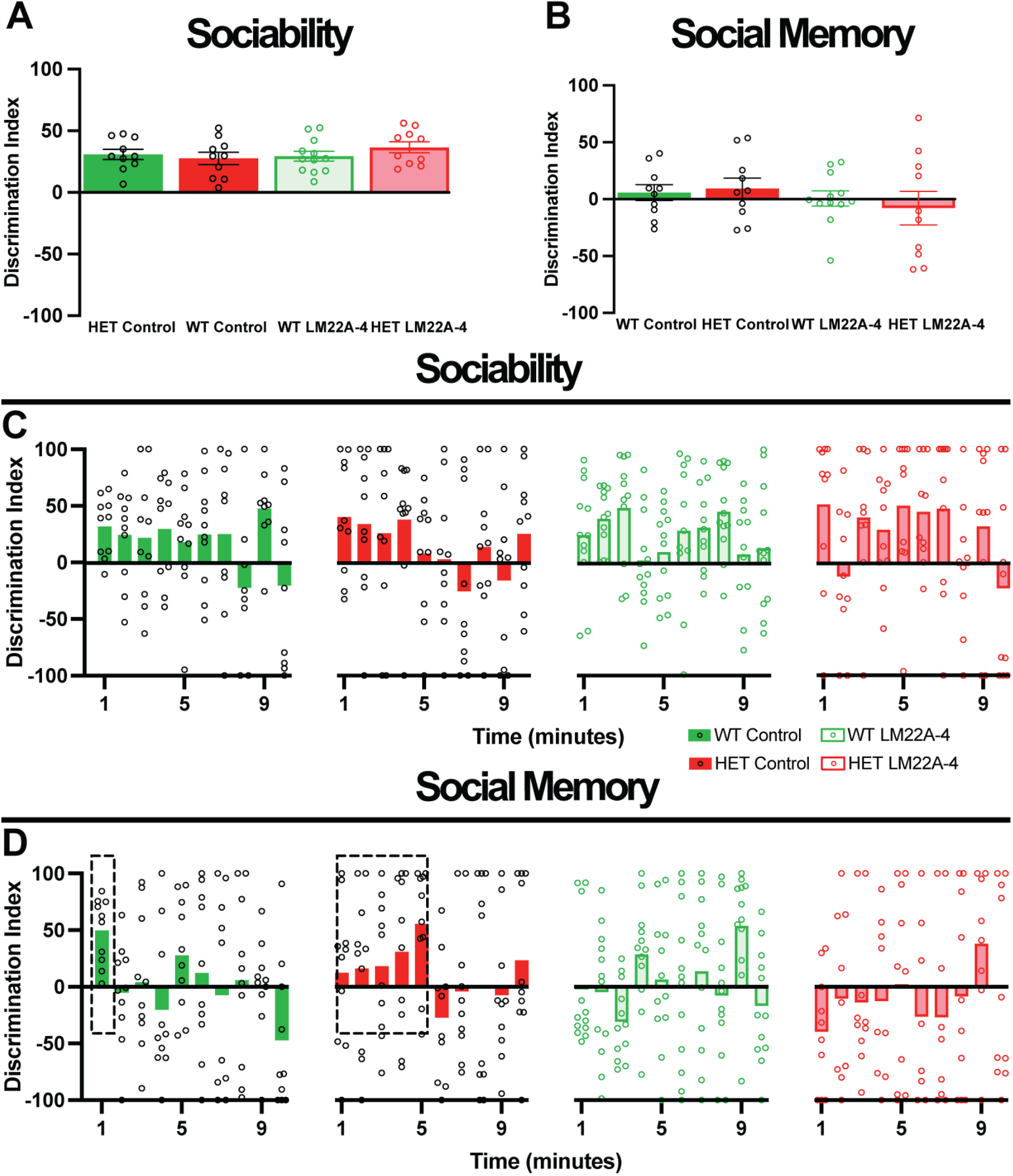
Female *Mecp2* HET mice have impaired social memory during the first minute of interaction with a novel mouse, but LM22A-4 treatment did not improve it. **(A,B)** Bulk discrimination indices of sociability and memory tests of female *Mecp2* HET mice and WT control littermates, treated with LM22A-4 or control. **(C,D)** Minute-by-minute discrimination indices of sociability and memory tests. Data are mean ±SEM.

Collectively, these findings contribute to the growing body of evidence regarding non-cell autonomous consequences of MeCP2 loss in the ‘mosaic’ female *Mecp2* HET brain. Hippocampal pyramidal neurons that express *Mecp2* have larger dendritic spines than neighboring neurons lacking *Mecp2*, and LM22A-4 partially rescues this atypical phenotype to levels similar to those in female WT neurons. These observations resemble the non-cell autonomous consequences of *Mecp2* loss in the primary motor cortex of female *Mecp2* HET mice, where *Mecp2*-expressing neurons show a reduction of 41% in spine density compared to *Mecp2*-lacking neurons (Belichenko et al., 2009). Moreover, the size of the cell body of *Mecp2*-expressing neurons in female *Mecp2* HET mice is smaller than that of neurons in female WT mice (Ribeiro and MacDonald, 2020; Rietveld et al., 2015; Wither et al., 2013).

### Effect of LM22A-4 on social behaviors in female *Mecp2* heterozygous mice

Four and 6-month-old female *Mecp2* HET mice were used to test the effects of a 4-week treatment with LM22A-4 on behaviors relevant to Rett syndrome. The total walking times during the 10-minutes of the standard 3-chamber test and novel unrestricted assay were not different between genotypes and were not affected by either LM22A-4 or saline treatments (p>0.9999, one-way ANOVA-Bonferroni’s; between all groups; Figure 4), indicating the lack of locomotion deficits at this age that could potentially confound the interpretations of other behavioral differences.

#### Standard 3-chamber test

We used the standard 3-chamber social interaction test (Moy et al., 2004) in 7-month-old mice, first for social preference (novel mouse vs. empty inverted pencil cup) and then for social memory (preference for prior novel mouse vs. new novel mouse), as described (Phillips et al., 2019). Using the average interaction time from the whole 10-min session, we found no differences between the discrimination indices for sociability (p>0.9999, ANOVA-Bonferroni’s; control WT n=10 mice vs. control *Mecp2* HET n=10; Figure 2A) and social memory (p>0.9999, ANOVA-Bonferroni’s; control WT vs. control *Mecp2* HET; Figure 2B). Consistent with this lack of differences, a 4-week treatment with LM22A-4 did not affect any of these measures neither in *Mecp2* HET nor in WT mice (p>0.9999, ANOVA-Bonferroni’s; LM22A-4 *Mecp2* HET n=10 vs. control *Mecp2* HET vs. LM22A-4 WT n=12 vs. control WT; Figure 2A,B).

Because female mice interact in shorter bouts with their preferred choice mainly at the start of the social preference test compared to males (Netser et al., 2017), the average interaction time from the whole 10-minute session may not be the most accurate measure of their behavior. Thus, we compared their discrimination index on a minute-by-minute basis throughout the 10-minute session. There were no differences in the sociability trial between female *Mecp2* HET and WT mice (p>0.9999, ANOVA-Bonferroni’s; control *Mecp2* HET vs. control WT; Figure 2C). However, during the social memory trial, most female WT mice (10 of 10 mice) had a positive discrimination index within the first minute of the test (boxed data in Figure 2D), suggesting they could quickly discriminate between a familiar mouse and a novel mouse. On the contrary, most female *Mecp2* HET mice (8 of 10 mice) took 5 minutes to reach a discrimination index similar to that of female WT mice (boxed data in Figure 2D; p>0.9999, ANOVA-Bonferroni’s; control *Mecp2* HET vs. control WT), indicating at least a delayed social memory. After those initial times, neither WT nor *Mecp2* HET mice showed clear socially motivated behaviors (p>0.9999, ANOVA-Bonferroni’s; control *Mecp2* HET vs. control WT; Figure 2D). Furthermore, LM22A-4 had no effects on these dynamics of social interactions, neither during the sociability test nor the social memory test (p>0.9999, ANOVA-Bonferroni’s; between all groups; Figure 2A-D). These observations are consistent with previous reports of female mice engaging differently than males during social interactions, where males engage in longer interaction bouts (Karlsson et al., 2015; Li and Dulac, 2018; Netser et al., 2017; Rodriguez et al., 2023; van den Berg et al., 2015; Williamson et al., 2019).

#### Unrestricted assay

Because the novel and familiar mice used as social stimuli are restrained under pencil cups during the standard social interaction test (Moy et al., 2004), which may result in experimental confounds (e.g. stress vocalizations by stimulus mice), we performed an unbiased assay of naturalistic behaviors shown by a test mouse when interacting with a novel mouse and a littermate, all freely moving in a large arena (12in x 16in) for 10 minutes. The 3 mice were videotaped from above, and their individual trajectories were tracked with *Motr* (Ohayon et al., 2013), which was used to train the machine-learning model *JAABA* (Kabra et al., 2013) with specific behaviors, thus producing an unbiased scoring of behaviors by the test mouse when interacting with freely moving novel and familiar mice. We successfully used this approach to reveal altered social behaviors in symptomatic male *Mecp2* KO mice (Phillips et al., 2019).

#### Non-aggressive (‘benign’) social interactions

Five-month-old female *Mecp2* HET mice (n=6 mice) showed more ‘face following’ (p=0.00358) and more ‘sniff nose’ (p=0.0005) than WT mice (n=6, ANOVA-Bonferroni’s; Figure 4A). A 4-week treatment with LM22A-4 in these mice reduced ‘sniff nose’ behavior in *Mecp2* HET mice (n=6) to levels comparable to those shown by WT mice (p=0.0487, ANOVA-Bonferroni’s; control *Mecp2* HET vs. LM22A-4 *Mecp2* HET) while having no effects in WT mice (n=9; Figure 4A).

In a slightly older cohort of 7-month-old female mice, there were no longer differences in ‘face following’ and ‘sniff nose’ (p>0.9999, ANOVA-Bonferroni’s, n=12 *Mecp2* HET mice, n=12 WT; Figure 4B). Furthermore, two other behaviors altered in male *Mecp2* KO mice, ‘rear sniffing’ and atypical ‘piggy-back jumping’ (Phillips et al., 2019) were unchanged between female *Mecp2* HET and WT mice (p>0.9999, ANOVA-Bonferroni’s; Figure 4B). The same results were obtained using the average discrimination indices of the whole interaction time or broken down in a minute-by-minute basis (p>0.9999, ANOVA-Bonferroni’s; between all groups; ‘face following’ Figure 3A,B; ‘rear sniffing’ Figure 3C,D; atypical ‘piggy-back jumping’ Figure 3E,F; ‘sniff nose’ Figure 4B). Consistent with this lack of differences between genotypes, LM22A-4 had no effects on ‘face following’, ’rear sniffing’, atypical ‘piggy-back jumping’, and ‘sniff nose’ behaviors shown by *Mecp2* HET (n=12 mice) and WT mice (n=12 mice), either using the 10-min average discrimination indices or in a minute-by-minute basis (p>0. 9999, ANOVA-Bonferroni’s; between all groups; ‘face following’ Figure 3A,B; ‘rear sniffing’ Figure 3C,D; atypical ‘piggy-back jumping’ Figure 3E,F; ‘sniff nose’ Figure 4B).

**Figure 3:**
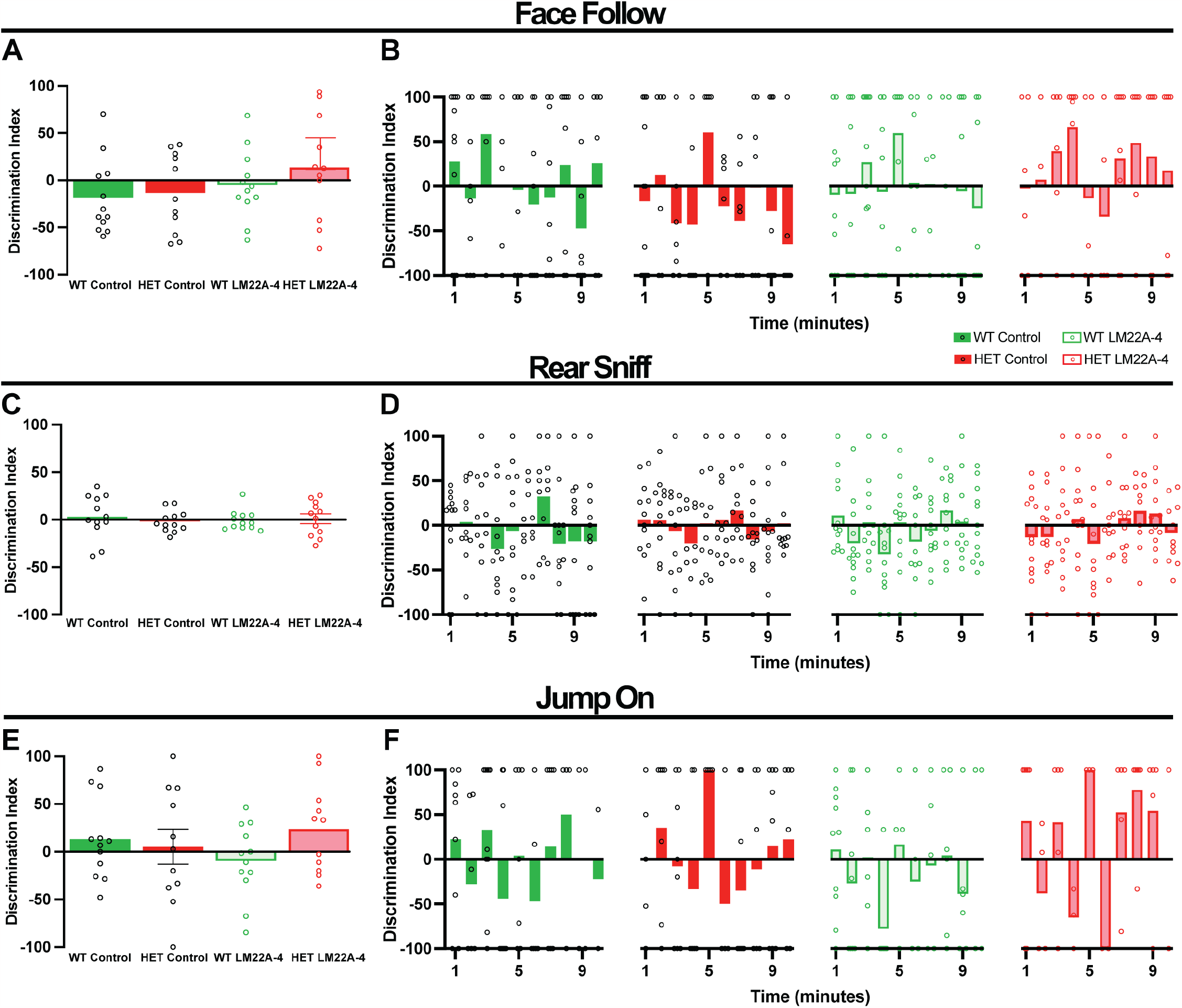
Social interactions in an unrestricted social assay were not altered in 7 months-old female *Mecp2* HET mice, and LM22A-4 treatment had no effect on them. (A) Bulk discrimination indices of ‘face follow’ behavior. (B) Minute-by-minute discrimination indices of ‘face follow’ behavior. (C) Bulk discrimination indices of ‘rear sniff’ behavior. (D) Minute-by-minute discrimination indices of ‘rear sniff’ behavior. (E) Bulk discrimination indices of ‘piggy-back jumping’ behavior. (F) Minute-by-minute discrimination indices of ‘piggy-back jumping’ behavior. Data are mean ±SEM.

**Figure 4:**
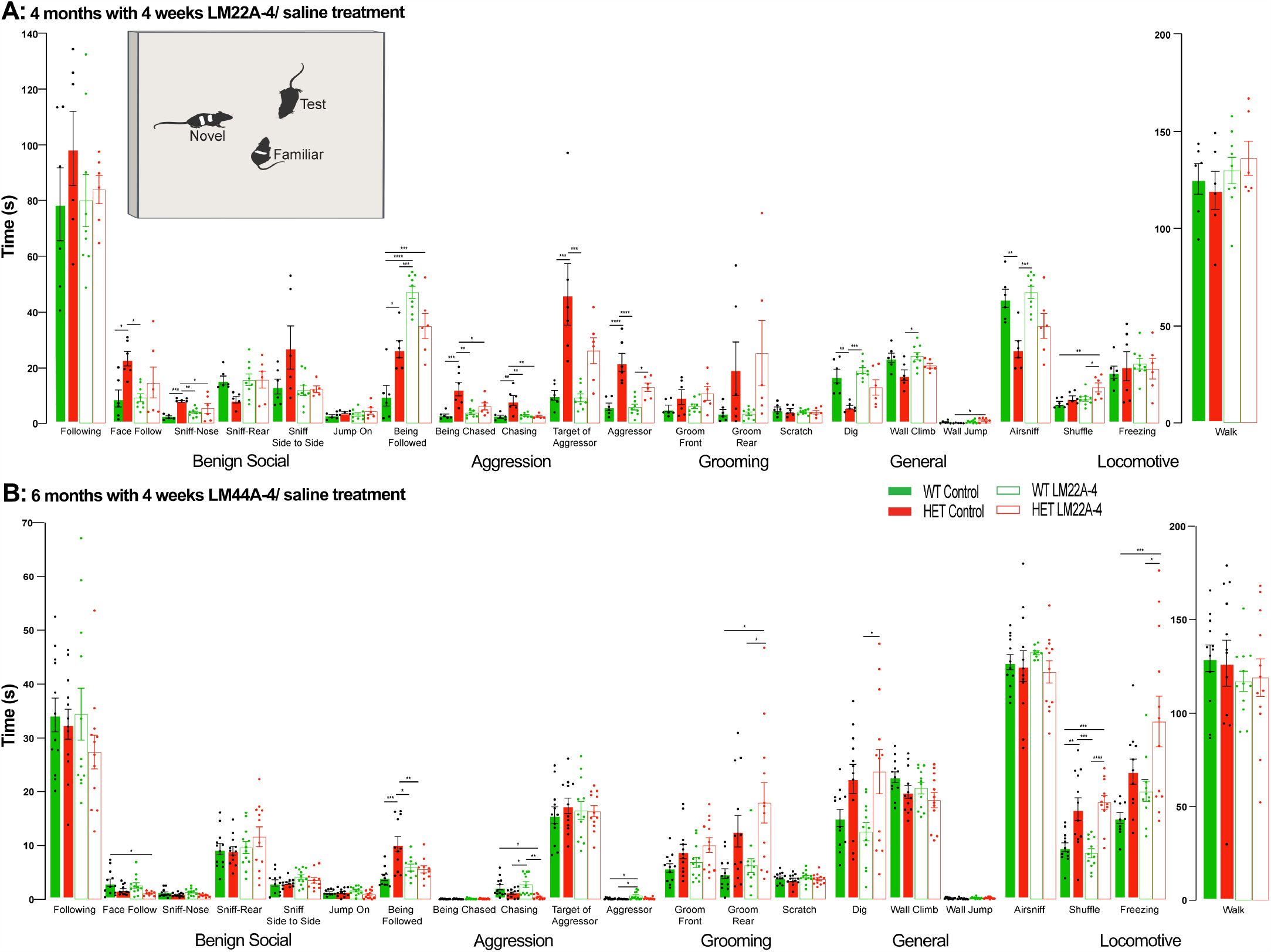
Unrestricted behavior analysis reveals enhanced aggressive behaviors in *Mecp2* HET mice, which LM22A-4 attenuates. **(Top inset)** Representative schematic of the unrestricted social assay. Time spent performing different behaviors using machine-learning scoring of social interactions between 3 unrestricted female mice: 1 *Mecp2* HET or WT with 1 novel and 1 familiar female WT. (A) Four months-old mice after a 4-week treatment with either LM22A-4 or saline. (B) Six months-old mice after a 4-week treatment with either LM22A-4 or saline. Data are mean ±SEM. *P<0.05, ** P<0.01, *** P<0.001, **** P<0.0001.

#### Target of social interactions

Five-month-old *Mecp2* HET mice were ‘followed’ more than WT mice (p=0.0130, ANOVA-Bonferroni’s; control WT vs. control *Mecp2* HET; Figure 4A), which was also observed in 6-month-old mice (p=0.0003, ANOVA-Bonferroni’s; control WT vs. control *Mecp2* HET; Figure 4B). Unexpectedly, 6-month-old mice *Mecp2* HET mice treated with LM22A-4 showed lower times being ‘followed’ than control *Mecp2* HET mice (p=0.0074, ANOVA-Bonferroni’s; LM22A-4 *Mecp2* HET vs. control *Mecp2* HET; Figure 4B).

#### Aggressive behaviors

Five-month-old female *Mecp2* HET mice exhibited longer epochs of ‘aggressive’ behaviors (p<0.0001) and ‘chasing’ behaviors (p=0.0008) than age-matched WT mice (ANOVA-Bonferroni; control *Mecp2* HET vs. control WT; Figure 4A). A 4-week treatment with LM22A-4 reduced the ‘chasing’ behaviors in *Mecp2* HET mice to WT levels (p=0.0033), but it did not affect ‘aggressive’ behaviors (p=0.0180, ANOVA-Bonferroni; LM22A-4 *Mecp2* HET vs. control *Mecp2* HET; Figure 4A). On the other hand, no differences between genotypes nor effects of LM22A-4 on ‘aggressive’ or ‘chasing’ behaviors were observed in 7-month-old mice (p>0.05, ANOVA-Bonferroni’s; Figure 4B).

#### Target of aggression

Five-month-old female *Mecp2* HET mice were the target of aggressive ‘chasing’ (p=0.0008) and ‘aggressive’ behaviors (p=0.0010) for longer times than age-matched WT mice (ANOVA-Bonferroni’s; control *Mecp2* HET control vs. control WT; Figure 4A). Interestingly, the 4-week LM22A-4 treatment reduced the duration in which *Mecp2* HET mice were being ‘chased’ aggressively (p=0.0359, ANOVA-Bonferroni’s; LM22A-4 *Mecp2* HET vs. control *Mecp2* HET; Figure 4A). However, LM22A-4 did not affect the duration of *Mecp2* HET mice being the target of ‘aggressive’ behaviors (p=0.1242, ANOVA-Bonferroni’s; LM22A-4 *Mecp2* HET vs. control *Mecp2* HET; Figure 4A).

#### Locomotion

Seven-month-old female *Mecp2* HET mice showed more ‘shuffling’ locomotion than age-matched WT mice (p= 0.0035, ANOVA-Bonferroni’s; control *Mecp2* HET vs. control WT; Figure 4B), but LM22A-4 did not affect it (p<0.0001, ANOVA-Bonferroni’s; LM22A-4 *Mecp2* HET vs. control *Mecp2* HET; Figure 4B).

#### General behaviors

Five-month-old female *Mecp2* HET mice showed shorter epochs of ‘digging’ (p=0.0023) and ‘air-sniffing’ behaviors (p=0.0052) than WT mice (ANOVA-Bonferroni’s; control *Mecp2* HET vs. control WT; Figure 4A), but LM22A-4 did not affect them (p>0.9999, ANOVA-Bonferroni’s; LM22A-4 *Mecp2* HET vs. control *Mecp2* HET; Figure 4B).

Taken together, these data indicate that social and aggression behaviors may be altered in female *Mecp2* HET mice prior to ‘shuffling’ during locomotion, and that LM22A-4 only improves some of the affected behaviors. Interestingly, social aggression was reported in male mice lacking *Mecp2* in either *Sim1*-expressing or serotonergic PET1-expressing neurons (Samaco et al., 2009). Similarly, *Mecp2* deletion in PET1-expressing neurons led to an aggressive phenotype in male mice, despite a lack of an anxiety phenotype, as tested in the ‘open field’ or ‘light-dark box’ assays (Fyffe et al., 2008). In a different study, male *Mecp2* KO mice showed hyper-reactive escape and defensive behaviors in a ‘mouse defense test battery’ assay (‘predator avoidance,’ ‘chase/flight’, ’closed door approach’, and ‘forced contact’ tests), despite the lack of an anxiety phenotype, as tested in the ‘elevated plus’ maze and ‘elevated zero’ maze (Pearson et al., 2015). Intriguingly, a phenotype-based genetic association study revealed that MeCP2 protein levels in mice and *MECP2* single nucleotide polymorphisms in schizophrenic individuals are associated with social aggression behaviors (Tantra et al., 2014). In addition, female *Mecp2* HET mice breed into 2 different genetic backgrounds, FVB/N x 129S6/SvEv and 129S6/SvEv x C57BL/6, showed altered sociability, contextual fear memory, and passive avoidance behaviors at 3 and 5 months of age, while at 4 months of age, these mice displayed reduced open field distance suggestive of a motor impairment (Samaco et al., 2013). Collectively, these data indicate that sex, genotype, and also possibly genetic background are highly determinant for the presentation of behavioral phenotypes, and therefore, of outcome measures and the efficacy of therapeutic approaches.

## Acknowledgements

We thank Ms. Lili Mao for mouse colony management and neuronal cultures, Dr. Takafumi Inoue (Waseda University, Tokyo, Japan) for data acquisition and analysis software, and Drs. Adrian Bird (University of Edinburg, Scotland) and Ben Philpot (University of North Carolina, USA) for MeCP2-GFP reporter mice. This work was supported by NIH grants HD-074418 (LP-M), MH-118563-01 (LP-M), T32-NS061788 (DM), MH-118563-04-S1 (DM), and the Rett Syndrome Research Trust (LP-M).

## Author Contributions

DM performed experiments, analyzed data, and wrote the manuscript; KA-B performed experiments and analyzed data; HE-B performed experiments and analyzed data; EM performed experiments and analyzed data; CC performed experiments and analyzed data; HR performed experiments and analyzed data; MLL analyzed data; TY provided unique reagents; FML provided unique reagents; WL performed experiments and analyzed data; LP-M designed experiments, analyzed data, and wrote the manuscript.

## MATERIAL AND METHODS

### Mice

Breeding pairs of mice lacking exon 3 of the X chromosome-linked *Mecp2* g e n e (B 6. C g - *Mecp2*tm1.1Jae, ‘Jaenisch’ strain in a pure C57BL/6 background) (Chen et al., 2001) were purchased from the Mutant Mouse Regional Resource Center at the University of California, Davis (stock #000415). A colony was established at The University of Alabama at Birmingham by mating male WT C57BL/6 mice with female *Mecp2* HET mice. Genotyping was performed by PCR of DNA samples from tail clips at weaning age P28. Male hemizygous *Mecp2* mice (i.e. *Mecp2* KO), develop typically until 5–6 weeks of age (P35-P42), when they begin to exhibit RTT-associated motor symptoms, such as hypoactivity, hind limb clasping, and reflex impairments (Chen et al., 2001). Male transgenic mice expressing GFP-tagged MeCP2 (Lyst et al., 2013); JAX stock #014610) were a gift from Drs. Adrian Bird (University of Edinburg, Scotland) and Ben Philpot (University of North Carolina, USA), and were crossed with female *Mecp2* HET mice, which allows identifying the cellular genotype in the female ‘mosaic’ brains resulting from X-chromosome inactivation. Female *Mecp2* HET mice develop similar RTT-associated symptoms than those observed in male *Mecp2* KO mice, but with a delayed progression, most being evident starting around 2-3 months of life (Samaco et al., 2013). Animals were handled and housed according to the Committee on Laboratory Animal Resources of the National Institutes of Health (NIH). All experimental protocols were annually reviewed and approved by the Institutional Animals Care and Use Committee (IACUC) of The University of Alabama at Birmingham.

### Organotypic slice cultures

Organotypic hippocampal slice cultures were prepared from male postnatal day 5-7 (P5-7) *Mecp2* KO mice and WT littermates as previously described (Chapleau et al., 2009; Pozzo Miller et al., 1993). Briefly, 500 µm-thick hippocampal slices were cut with a custom-made wire-slicer (Katz, 1987), plated on tissue culture plate inserts (Millicell-CM, Millipore) inside 6-well plates with culture medium containing 78% Neurobasal-A without phenol red (Invitrogen), 20% heat-inactivated equine serum (Invitrogen), 2% B27 supplement (Invitrogen), 0.5 mM L-glutamine (Invitrogen), and placed in an incubator at 36°C, 5% CO2, 90% relative humidity. Serum was titrated out over 3 days *in vitro* (DIV), and treatments were made in serum-free culture media, as described (Chapleau et al., 2008).

### Particle-mediated gene transfer

Seven DIV slice cultures were transfected with enhanced yellow fluorescent protein (eYFP) by particle-mediated gene transfer using a Helios Gene Gun (Bio-Rad), as described (Alonso et al., 2004; Chapleau et al., 2009). Briefly, the eYFP encoding cDNA plasmid (Clontech) was precipitated onto colloidal Au particles (1.6µm) and coated onto Tefzel tubing. Slices were bombarded with Au particles accelerated by ∼85 pounds per square inch (PSI) of He gas from a distance of 2 cm using a modified ‘gene-gun’ nozzle with a 2µm filter. Biolistic transfections were performed 24 hours after culture medium was changed to serum-free culture medium with antibiotics (1:100 penicillin/streptomycin, Invitrogen) and antimycotics (1:100 Fungizone, Invitrogen); 24 hours after transfection, the culture medium was changed to serum-free medium without antibiotics or antimycotics.

### *In vitro* BDNF and LM22A-4 treatment

Nine DIV hippocampal slice cultures were randomly assigned to treatment groups with the following drugs dissolved in serum-free medium: recombinant human BDNF (250 ng/mL, Promega); BDNF + K-252a (200 nM, Calbiochem); K-252a alone; LM22A-4 (500 nM); LM22A-4 + K-252a; LM22A-4 + ANA-12 (100 µM, Sigma); ANA-12 alone. Culture medium was removed from culture wells and replaced with drug-containing medium; in addition, 50 µL of drug-containing medium was gently applied on top of each slice. All treatments lasted 48 hours. Eleven DIV slice cultures were fixed with 4% paraformaldehyde in 100 mM phosphate buffer (PB) overnight at 4°C, washed with 100 mM phosphate-buffered saline (PBS), trimmed from the filter membrane of the cultured inserts, and mounted on glass slides with Vectashield (Ve c t o r Laboratories).

### *In vivo* LM22A-4 treatment

Four to 6 months-old female *Mecp2* HET mice and female MeCP2-GFP HET mice, and their age-matched female WT littermates received intraperitoneal (i.p.) injections of either sterile LM22A-4 (50 mg/kg) or vehicle (0.9% NaCl) twice daily for 4 weeks, following an established dosing regime with efficacy in other studies using *Mecp2* mutant mice (Kron et al., 2012; Li et al., 2017; Schmid et al., 2012). Mice were randomly assigned to each treatment; LM22A-4 was prepared daily from sterile stocks and dissolved in sterile saline.

### Intracellular loading of fluorescent dye in *ex vivo* hippocampal slices

Mice were deeply anesthetized with a mixture of 100mg/kg ketamine and 10mg/kg xylazine, and transcardially perfused with ice-cold ‘cutting’ artificial cerebrospinal fluid (aCSF), containing (in mM): 87 NaCl, 2.5 KCl, 0.5 CaCl2, 7 MgCl2, 1.25 NaH2PO4, 25 NaHCO3, 25 glucose, and 75 sucrose, bubbled with 95% O2/5% CO2. The brain was rapidly removed, and cut transversely at 300µm using a vibrating blade microtome (VT1200S, Leica Microsystems) in the same ice-cold ‘cutting’ aCSF. Slices were transferred to normal oxygenated Acsf containing (in mM): 119 NaCl, 2.5 KCl, 2.5 CaCl2, 1.3 MgCl2, 1.3 NaH2PO4, 26 NaHCO3, and 20 glucose (95% O2/5% CO2), at 32°C for 30 min, and then allowed to recover for 1 hour at room temperature before use. Individual slices were transferred to a submerged chamber mounted on a fixed-stage upright microscope (Axioskop FS or AxioExaminer D1, Zeiss) and continuously perfused at room temperature with normal oxygenated aCSF containing (in mM): 119 NaCl, 2.5 KCl, 2.5 CaCl2, 1.3 MgCl2, 1.3 NaH2PO4, 26 NaHCO3, and 20 glucose (95% O2/5% CO2). Pyramidal neurons in CA1 stratum pyramidale were visualized by infrared differential interference contrast microscopy with water-immersion objectives (63X 0.9 NA, or 20X 1.0NA plus 0.50-4X zoom, Zeiss); in MeCP2-GFP mice, GFP was imaged with 475nm LED illumination (X-Cite Turbo, Excelitas), a GFP cube (Semrock), and a QuantEM:512SC cooled CCD (Photometrics). Whole-cell pipettes contained (in mM): 120 Cs-gluconate, 17.5 CsCl, 10 Na-HEPES, 4 Mg-ATP, 0.4 Na-GTP, 10 Na2-creatine phosphate, 0.2 Na-EGTA, and biocytin (8mM, Sigma Aldrich), 290-300mOsm, pH 7.3 (final resistance, 3-4MΩ). After ∼15min of whole-cell access to allow biocytin loading, slices were fixed in 4% paraformaldehyde in 100mM PBS, and stained with streptavidin-conjugated Alexa Fluor-488 (Life Technologies), as described (Li et al., 2016).

### Confocal microscopy and dendritic spine analyses

eYFP-expressing CA1 and CA3 pyramidal neurons in hippocampal slice cultures from male *Mecp2* KO mice and male WT littermates, and biocytin-filled, Alexa Fluor-488 labeled CA1 pyramidal neurons in hippocampal slices from female MeCP2-GFP HET mice and female WT littermates were selected for confocal imaging if they show fluorescent label throughout the entire dendritic tree and lacked signs of degeneration (e.g. dendritic ‘blebbing’). High-resolution z-stack images of apical secondary and tertiary dendrites were acquired with oil-immersion 60X (NA 1.42) objectives (PlanApo) plus 3X digital zoom in either a Fluoview FV-300 (Olympus) or a LSM-800 (Zeiss) confocal microscope; image stacks were acquired at 0.1 µm intervals in the z-plane. Z-stack image stacks were 3D-reconstructed, surface rendered, and analyzed semi-automatically with the Filament Tracing module of Imaris (Bitplane) (Swanger et al., 2011). Image stacks were loaded into Imaris, a region-of-interest (ROI) including a 40-80µm-long dendritic shaft was selected, and Filament Tracing was used to trace and render dendrites and their associated spines. Spines were defined as dendritic protrusions shorter than 3µm. The numerical densities of spines were calculated for each dendritic segment, and normalized to 10µm of dendritic length. The relative intensity of individual spines and their parent dendritic shafts was used to estimate the volume of individual spines (Benavides-Piccione et al., 2013). An experimenter blind to genotype and treatment performed dendritic spine analyses.

### Behavioral analyses

All handling and testing were done in the dark phase of the standard 12 hr light/12 hr dark cycle (6am ON, 6pm OFF), with the experimenter wearing a red headlamp and infrared illumination for digital videography.

#### Standard 3-chamber social test

We used the standard 3-chamber social interaction test (Moy et al., 2004). Prior to testing, mice underwent a 3-day acclimation period during which they were familiarized to the experimenter with handling for 3 minutes each day at the same time as testing. The testing environment was a 3-chambered box which contained 2 empty inverted pencil cups placed in the 2 side chambers. Mice were placed in the center chamber and allowed to freely explore for 5 minutes. After this initial acclimation period, mice were directed to return to the center chamber, and barriers were positioned over the side openings. At this time, a novel mouse was placed beneath one of the previously empty pencil cups in one of the 2 side chambers, alternating between trials. The barriers were then removed, and the test mouse was allowed to freely explore the chambers for 10 minutes. Subsequently, the test mouse was again directed to return to the center chamber, and barriers were positioned over the side openings. A second novel mouse was placed beneath the formerly empty pencil cup, the previously added novel mouse is now considered familiar. After removing the side barriers, the test mouse was allowed to explore the chambers for an additional 10 minutes. Following each test, the apparatus was meticulously cleaned with 70% isopropanol. The time in which the test mouse spent actively sniffing each pencil cup, containing either a social conspecific or empty, was quantified. For the sociability test, a discrimination index was calculated as [(Time investigating Mouse Cup - Time investigating Empty Cup) / (Time investigating Mouse Cup + Time investigating Empty Cup) * 100]. For the social memory trial, the discrimination index was calculated as [(Time investigating Novel Mouse Cup - (Time investigating Familiar Mouse Cup) / (Time investigating Novel Mouse Cup + (Time investigating Familiar Mouse Cup) *100]. Test mice that spent over 75% of the acclimation time in a single compartment were excluded from the study. Statistical analyses were performed on the discrimination indices for each test mouse.

#### Unrestricted assay

One week prior to testing, the back of sentinel novel and familiar mice was dyed with blond hair dye (Born Blonde Maxi, Clairol) with different patterns for computer vision tracking. Mice were habituated to the testing arena, an open field box (12in x 16in) containing clean bedding, for 3 days prior to the testing day, 10 minutes per day. During the testing day, test mice were habituated to the testing arena for 10 minutes. Sentinel mice, including one cage-mate and one novel mouse from a different cage, were placed in the testing arena and were allowed to interact freely for 10 minutes. After this time, sentinel mice were placed into a neutral cage, and the test mouse was returned to their home cage. The arena was meticulously cleaned with 70% isopropanol and filled with new bedding between each test. Sentinel mice interacted with a maximum of 5 test mice and were no longer used within the study if they fought with other sentinels or displayed excessive grooming phenotypes. Test videos were loading into Motr (github.com/motr/motr) (Ohayon et al., 2013) to create tracks that were exported to the machine-learning model Janelia Automatic Animal Behavior Annotator (JAABA; gi thub. com/ kristinbranson/JAABA) (Kabra et al., 2013) for unbiased behavioral scoring. JAABA classifiers were first trained on pilot data sets. Behavioral scores for social memory and other behaviors were taken from the entire video, as different behaviors emerged at later times during the 10min trial, and social behavior times were pooled between novel and cage-mate mouse. Mice not interacting with sentinels for more than 3 sec (out of 240 sec) were excluded from analyses. Statistical tests were performed on the time per behavior for each test mouse.

### Statistical analyses

All experiments were repeated in at least 3 independent slice cultures or *ex vivo* acute slice preparations. All statistical analyses were performed using Prism (GraphPad Software), P <0.05 was considered significant. Data are presented as mean ± standard error of the mean, and were compared using unpaired Student’s t-tests for 2 groups or one-way ANOVA with Bonferroni’s post-hoc test for more than 2 groups. Data with more than 3 groups that did not fit the normal distribution were analyzed using the Kruskal-Wallis test, with Dunn’s post-hoc test for multiple comparisons; post-hoc tests were only performed for ANOVA and Kruskal-Wallis tests that reported P < 0.05. The Kolmogorov-Smirnov (K-S) test was used for comparisons of cumulative frequency distributions.

## SUPPLEMENTAL FIGURES

**Supplemental Figure 1:**
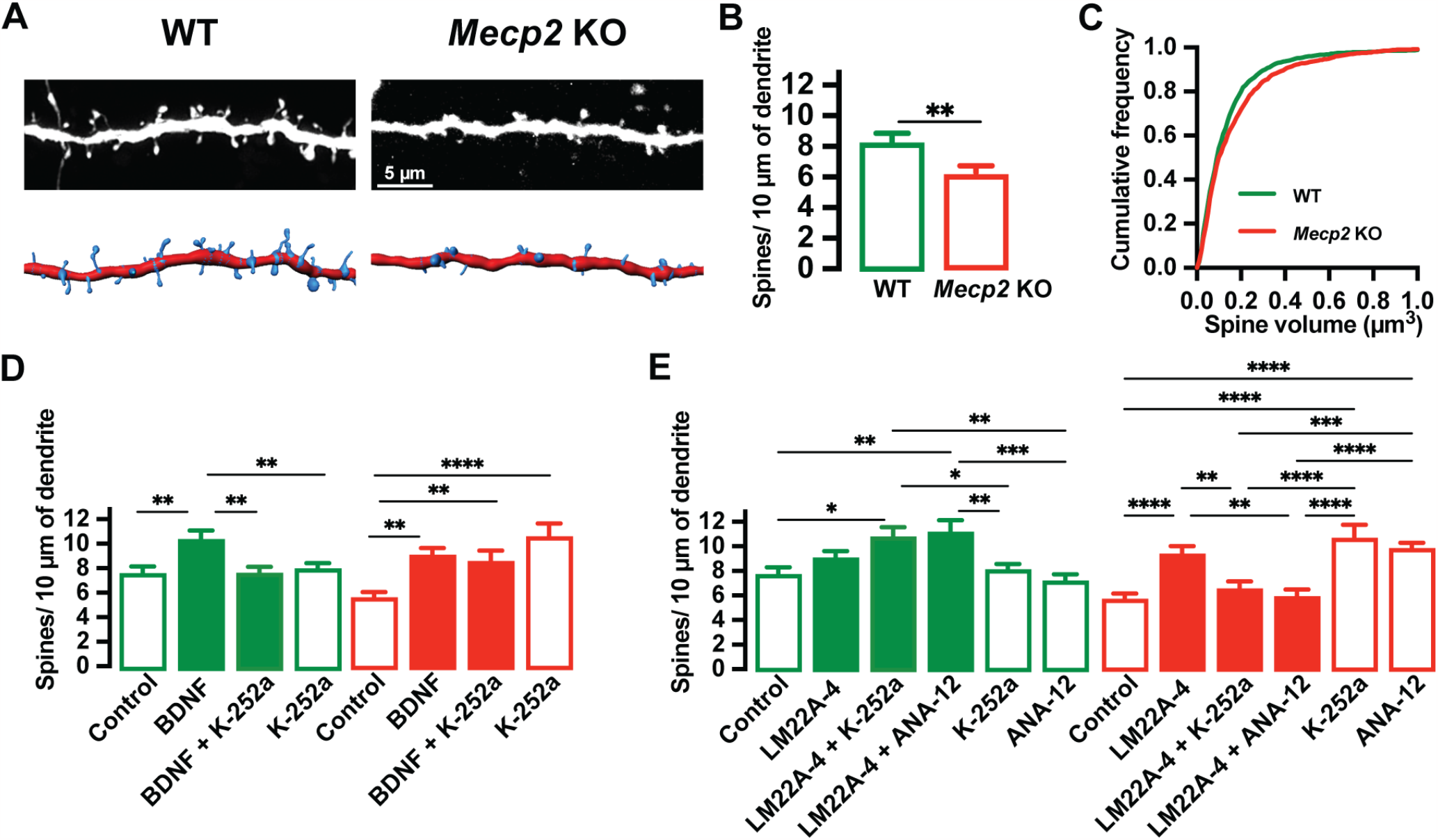
LM22A-4 increases dendritic spine density in pyramidal neurons of hippocampal slice cultures from male *Mecp2* KO mice via activation of TrkB receptors. (A) Representative images of eYFP-expressing pyramidal neurons in cultured hippocampal slices from P7 male *Mecp2* KO mice. (B) Spine density per µm of dendrite in WT and *Mecp2* KO mice. (C) Cumulative frequency of spine volume in WT and *Mecp2* KO neurons. (D) Spine density per µm of dendrite in WT and *Mecp2* KO mice following treatment of BDNF and inhibitor K-252a. (E) Spine density per µm of dendrite in WT and *Mecp2* KO mice following treatment of LM22A-4 and inhibitors K-252a and ANA-12. *P<0.05, ** P<0.01, *** P<0.001, **** P<0.0001.

**Supplemental Figure 2:**
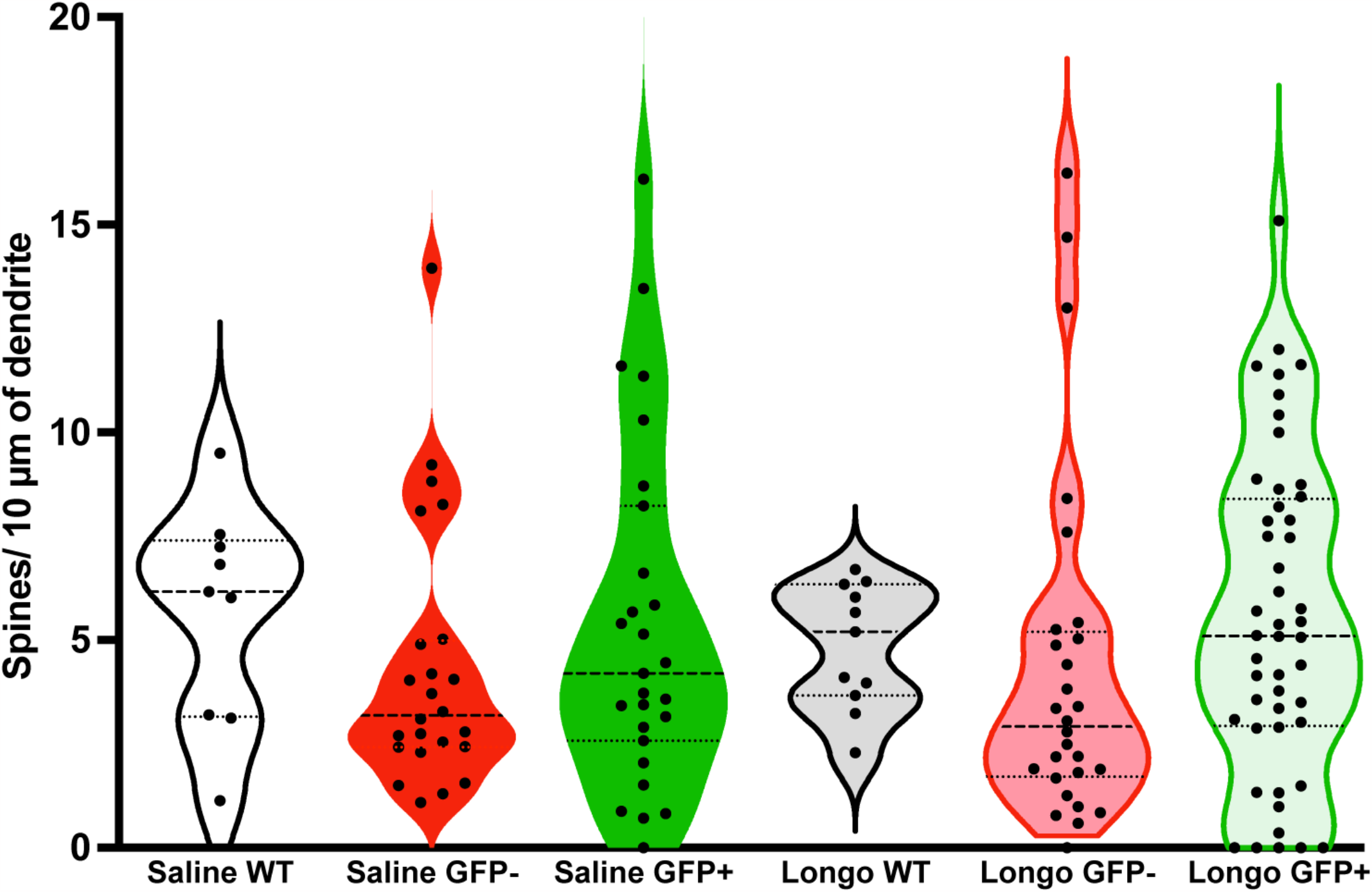
Dendritic spine density is similar in the two cellular genotypes of female MeCP2-GFP HET mice, and comparable to that in female WT mice. Spine density per µm of dendrite in WT and MeCP2-HET-GFP expressing or lacking neurons treated with control or LM22A-4. Data are mean ±SEM.

## REFERENCES

Abuhatzira, L., Makedonski, K., Kaufman, Y., Razin, A. and Shemer, R. (2007). MeCP2 deficiency in the brain decreases BDNF levels by REST/ CoREST-mediated repression and increases TRKB production. Epigenetics 2, 214–22.

Adams, I., Yang, T., Longo, F. M. and Katz, D. M. (2020). Restoration of motor learning in a mouse model of Rett syndrome following long-term treatment with a novel small-molecule activator of TrkB. Dis Model Mech 13.

Alonso, M., Medina, J. H. and Pozzo-Miller, L. (2004). ERK1/2 activation is necessary for BDNF to increase dendritic spine density in hippocampal CA1 pyramidal neurons. Learn Mem 11, 172–8.

Amir, R. E., Van den Veyver, I. B., Wan, M., Tran, C. Q., Francke, U. and Zoghbi, H. Y. (1999). Rett syndrome is caused by mutations in X-linked MECP2, encoding methyl-CpG-binding protein 2. Nat Genet 23, 185–8.

Belichenko, N. P., Belichenko, P. V. and Mobley, W. C. (2009). Evidence for both neuronal cell autonomous and nonautonomous effects of methyl-CpG-binding protein 2 in the cerebral cortex of female mice with Mecp2 mutation. Neurobiol Dis 34, 71–7.

Benavides-Piccione, R., Fernaud-Espinosa, I., Robles, V., Yuste, R. and DeFelipe, J. (2013). Age-based comparison of human dendritic spine structure using complete three-dimensional reconstructions. Cereb Cortex 23, 1798–810.

Calfa, G., Hablitz, J. J. and Pozzo-Miller, L. (2011). Network hyperexcitability in hippocampal slices from Mecp2 mutant mice revealed by voltage-sensitive dye imaging. J Neurophysiol 105, 1768–84.

Calfa, G., Li, W., Rutherford, J. M. and Pozzo-Miller, L. (2015). Excitation/inhibition imbalance and impaired synaptic inhibition in hippocampal area CA3 of Mecp2 knockout mice. Hippocampus 25, 159–68.

Cazorla, M., Premont, J., Mann, A., Girard, N., Kellendonk, C. and Rognan, D. (2011). Identification of a low-molecular weight TrkB antagonist with anxiolytic and antidepressant activity in mice. J Clin Invest 121, 1846–57.

Chahrour, M., Jung, S. Y., Shaw, C., Zhou, X., Wong, S. T., Qin, J. and Zoghbi, H. Y. (2008). MeCP2, a key contributor to neurological disease, activates and represses transcription. Science 320, 1224–9.

Chang, Q., Khare, G., Dani, V., Nelson, S. and Jaenisch, R. (2006). The disease progression of Mecp2 mutant mice is affected by the level of BDNF expression. Neuron 49, 341–8.

Chapleau, C. A., Boggio, E. M., Calfa, G., Percy, A. K., Giustetto, M. and Pozzo-Miller, L. (2012). Hippocampal CA1 pyramidal neurons of Mecp2 mutant mice show a dendritic spine phenotype only in the presymptomatic stage. Neural Plast 2012, 976164.

Chapleau, C. A., Calfa, G. D., Lane, M. C., Albertson, A. J., Larimore, J. L., Kudo, S., Armstrong, D. L., Percy, A. K. and Pozzo-Miller, L. (2009). Dendritic spine pathologies in hippocampal pyramidal neurons from Rett syndrome brain and after expression of Rett-associated MECP2 mutations. Neurobiol Dis 35, 219–33.

Chapleau, C. A., Carlo, M. E., Larimore, J. L. and Pozzo-Miller, L. (2008). The actions of BDNF on dendritic spine density and morphology in organotypic slice cultures depend on the presence of serum in culture media. J Neurosci Methods 169, 182–90.

Chapleau, C. A. and Pozzo-Miller, L. (2012). Divergent roles of p75NTR and Trk receptors in BDNF’s effects on dendritic spine density and morphology. Neural Plast 2012, 578057.

Chen, R. Z., Akbarian, S., Tudor, M. and Jaenisch, R. (2001). Deficiency of methyl-CpG binding protein-2 in CNS neurons results in a Rett-like phenotype in mice. Nat Genet 27, 327–31.

Chen, W. G., Chang, Q., Lin, Y., Meissner, A., West, A. E., Griffith, E. C., Jaenisch, R. and Greenberg, M. E. (2003). Derepression of BDNF transcription involves calcium-dependent phosphorylation of MeCP2. Science 302, 885–9.

Fyffe, S. L., Neul, J. L., Samaco, R. C., Chao, H. T., Ben-Shachar, S., Moretti, P., McGill, B. E., Goulding, E. H., Sullivan, E., Tecott, L. H. et al. (2008). Deletion of Mecp2 in Sim1-expressing neurons reveals a critical role for MeCP2 in feeding behavior, aggression, and the response to stress. Neuron 59, 947–58.

Geraghty, A. C., Gibson, E. M., Ghanem, R. A., Greene, J. J., Ocampo, A., Goldstein, A. K., Ni, L., Yang, T., Marton, R. M., Pasca, S. P. et al. (2019). Loss of Adaptive Myelination Contributes to Methotrexate Chemotherapy-Related Cognitive Impairment. Neuron 103, 250–265 e8.

Gines, S., Paoletti, P. and Alberch, J. (2010). Impaired TrkB-mediated ERK1/2 activation in huntington disease knock-in striatal cells involves reduced p52/p46 Shc expression. J Biol Chem 285, 21537–48.

Gu, F., Parada, I., Yang, T., Longo, F. M. and Prince, D. A. (2022). Chronic partial TrkB activation reduces seizures and mortality in a mouse model of Dravet syndrome. Proc Natl Acad Sci U S A 119.

Hartmann, D., Drummond, J., Handberg, E., Ewell, S. and Pozzo-Miller, L. (2012). Multiple approaches to investigate the transport and activity-dependent release of BDNF and their application in neurogenetic disorders. Neural Plast 2012, 203734.

Kabra, M., Robie, A. A., Rivera-Alba, M., Branson, S. and Branson, K. (2013). JAABA: interactive machine learning for automatic annotation of animal behavior. Nat Methods 10, 64–7.

Karlsson, S. A., Haziri, K., Hansson, E., Kettunen, P. and Westberg, L. (2015). Effects of sex and gonadectomy on social investigation and social recognition in mice. BMC Neurosci 16, 83.

Katz, D. M. (2014). Brain-derived neurotrophic factor and Rett syndrome. Handb Exp Pharmacol 220, 481–95.

Katz, D. M., Berger-Sweeney, J. E., Eubanks, J. H., Justice, M. J., Neul, J. L., Pozzo-Miller, L., Blue, M. E., Christian, D., Crawley, J. N., Giustetto, M. et al. (2012). Preclinical research in Rett syndrome: setting the foundation for translational success. Dis Model Mech 5, 733–45.

Katz, L. C. (1987). Local circuitry of identified projection neurons in cat visual cortex brain slices. J Neurosci 7, 1223–49.

Kron, M., Howell, C. J., Adams, I. T., Ransbottom, M., Christian, D., Ogier, M. and Katz, D. M. (2012). Brain activity mapping in Mecp2 mutant mice reveals functional deficits in forebrain circuits, including key nodes in the default mode network, that are reversed with ketamine treatment. J Neurosci 32, 13860–72.

Li, W., Bellot-Saez, A., Phillips, M. L., Yang, T., Longo, F. M. and Pozzo-Miller, L. (2017). A small-molecule TrkB ligand restores hippocampal synaptic plasticity and object location memory in Rett syndrome mice. Dis Model Mech 10, 837–845.

Li, W., Calfa, G., Larimore, J. and Pozzo-Miller, L. (2012). Activity-dependent BDNF release and TRPC signaling is impaired in hippocampal neurons of Mecp2 mutant mice. Proc Natl Acad Sci U S A 109, 17087–92.

Li, W. and Pozzo-Miller, L. (2014). BDNF deregulation in Rett syndrome. Neuropharmacology 76 Pt C, 737–46.

Li, W., Xu, X. and Pozzo-Miller, L. (2016). Excitatory synapses are stronger in the hippocampus of Rett syndrome mice due to altered synaptic trafficking of AMPA-type glutamate receptors. Proc Natl Acad Sci U S A 113, E1575–84.

Li, Y. and Dulac, C. (2018). Neural coding of sex-specific social information in the mouse brain. Curr Opin Neurobiol 53, 120–130.

Lyst, M. J., Ekiert, R., Ebert, D. H., Merusi, C., Nowak, J., Selfridge, J., Guy, J., Kastan, N. R., Robinson, N. D., de Lima Alves, F. et al. (2013). Rett syndrome mutations abolish the interaction of MeCP2 with the NCoR/SMRT co-repressor. Nat Neurosci 16, 898–902.

Martinowich, K., Hattori, D., Wu, H., Fouse, S., He, F., Hu, Y., Fan, G. and Sun, Y. E. (2003). DNA methylation-related chromatin remodeling in activity-dependent BDNF gene regulation. Science 302, 890–3.

Massa, S. M., Yang, T., Xie, Y., Shi, J., Bilgen, M., Joyce, J. N., Nehama, D., Rajadas, J. and Longo, F. M. (2010). Small molecule BDNF mimetics activate TrkB signaling and prevent neuronal degeneration in rodents. J Clin Invest 120, 1774–85.

Moy, S. S., Nadler, J. J., Perez, A., Barbaro, R. P., Johns, J. M., Magnuson, T. R., Piven, J. and Crawley, J. N. (2004). Sociability and preference for social novelty in five inbred strains: an approach to assess autistic-like behavior in mice. Genes Brain Behav 3, 287–302.

Nan, X., Campoy, F. J. and Bird, A. (1997). MeCP2 is a transcriptional repressor with abundant binding sites in genomic chromatin. Cell 88, 471–81.

Netser, S., Haskal, S., Magalnik, H. and Wagner, S. (2017). A novel system for tracking social preference dynamics in mice reveals sex- and strain-specific characteristics. Mol Autism 8, 53.

Neul, J. L. and Zoghbi, H. Y. (2004). Rett syndrome: a prototypical neurodevelopmental disorder. Neuroscientist 10, 118–28.

Ogier, M., Wang, H., Hong, E., Wang, Q., Greenberg, M. E. and Katz, D. M. (2007). Brain-derived neurotrophic factor expression and respiratory function improve after ampakine treatment in a mouse model of Rett syndrome. J Neurosci 27, 10912–7.

Ohayon, S., Avni, O., Taylor, A. L., Perona, P. and Roian Egnor, S. E. (2013). Automated multi-day tracking of marked mice for the analysis of social behaviour. J Neurosci Methods 219, 10–9.

Pearson, B. L., Defensor, E. B., Blanchard, D. C. and Blanchard, R. J. (2015). Applying the e t h o e x p e r i m e n t a l a p p r o a c h t o neurodevelopmental syndrome research reveals exaggerated defensive behavior in Mecp2 mutant mice. Physiol Behav 146, 98–104.

Percy, A. K. and Lane, J. B. (2005). Rett syndrome: model of neurodevelopmental disorders. J Child Neurol 20, 718–21.

Phillips, M. L., Robinson, H. A. and Pozzo-Miller, L. (2019). Ventral hippocampal projections to the medial prefrontal cortex regulate social memory. Elife 8.

Poduslo, J. F. and Curran, G. L. (1996). Permeability at the blood-brain and blood-nerve barriers of the neurotrophic factors: NGF, CNTF, NT-3, BDNF. Brain Res Mol Brain Res 36, 280–6.

Pozzo Miller, L. D., Petrozzino, J. J., Mahanty, N. K. and Connor, J. A. (1993). Optical imaging of cytosolic calcium, electrophysiology, and ultrastructure in pyramidal neurons of organotypic slice cultures from rat hippocampus. Neuroimage 1, 109–20.

Ribeiro, M. C. and MacDonald, J. L. (2020). Sex differences in Mecp2-mutant Rett syndrome model mice and the impact of cellular mosaicism in phenotype development. Brain Res 1729, 146644.

Rietveld, L., Stuss, D. P., McPhee, D. and Delaney, K. R. (2015). Genotype-specific effects of Mecp2 loss-of-function on morphology of Layer V pyramidal neurons in heterozygous female Rett syndrome model mice. Front Cell Neurosci 9, 145.

Rodriguez, L. A., Kim, S. H., Page, S. C., Nguyen, C. V., Pattie, E. A., Hallock, H. L., Valerino, J., Maynard, K. R., Jaffe, A. E. and Martinowich, K. (2023). The basolateral amygdala to lateral septum circuit is critical for regulating social novelty in mice. Neuropsychopharmacology 48, 529–539.

Samaco, R. C., Mandel-Brehm, C., Chao, H. T., Ward, C. S., Fyffe-Maricich, S. L., Ren, J., Hyland, K., Thaller, C., Maricich, S. M., Humphreys, P. et al. (2009). Loss of MeCP2 in aminergic neurons causes cell-autonomous defects in neurotransmitter synthesis and specific behavioral abnormalities. Proc Natl Acad Sci U S A 106, 21966–71.

Samaco, R. C., McGraw, C. M., Ward, C. S., Sun, Y., Neul, J. L. and Zoghbi, H. Y. (2013). Female Mecp2(+/-) mice display robust behavioral deficits on two different genetic backgrounds providing a framework for pre-clinical studies. Hum Mol Genet 22, 96–109.

Schmid, D. A., Yang, T., Ogier, M., Adams, I., Mirakhur, Y., Wang, Q., Massa, S. M., Longo, F. M. and Katz, D. M. (2012). A TrkB small molecule partial agonist rescues TrkB phosphorylation deficits and improves respiratory function in a mouse model of Rett syndrome. J Neurosci 32, 1803–10.

Segal, I., Korkotian, I. and Murphy, D. D. (2000). Dendritic spine formation and pruning: common cellular mechanisms? Trends Neurosci 23, 53–7.

Simmons, D. A., Belichenko, N. P., Yang, T., Condon, C., Monbureau, M., Shamloo, M., Jing, D., Massa, S. M. and Longo, F. M. (2013). A small molecule TrkB ligand reduces motor impairment and neuropathology in R6/2 and BACHD mouse models of Huntington’s disease. J Neurosci 33, 18712–27.

Swanger, S. A., Yao, X., Gross, C. and Bassell, G. J. (2011). Automated 4D analysis of dendritic spine morphology: applications to stimulus-induced spine remodeling and pharmacological rescue in a disease model. Mol Brain 4, 38.

Tantra, M., Hammer, C., Kastner, A., Dahm, L., Begemann, M., Bodda, C., Hammerschmidt, K., Giegling, I., Stepniak, B., Castillo Venzor, A. et al. (2014). Mild expression differences of MECP2 influencing aggressive social behavior. EMBO Mol Med 6, 662–84.

Tapia-Arancibia, L., Aliaga, E., Silhol, M. and Arancibia, S. (2008). New insights into brain BDNF function in normal aging and Alzheimer disease. Brain Res Rev 59, 201–20.

Tyler, W. J. and Pozzo-Miller, L. D. (2001). BDNF enhances quantal neurotransmitter release and increases the number of docked vesicles at the active zones of hippocampal excitatory synapses. J Neurosci 21, 4249–58.

van den Berg, W. E., Lamballais, S. and Kushner, S. A. (2015). Sex-specific mechanism of social hierarchy in mice. Neuropsychopharmacology 40, 1364–72.

Wang, H., Chan, S. A., Ogier, M., Hellard, D., Wang, Q., Smith, C. and Katz, D. M. (2006). Dysregulation of brain-derived neurotrophic factor expression and neurosecretory function in Mecp2 null mice. J Neurosci 26, 10911–5.

Williamson, C. M., Lee, W., DeCasien, A. R., Lanham, A., Romeo, R. D. and Curley, J. P. (2019). Social hierarchy position in female mice is associated with plasma corticosterone levels and hypothalamic gene expression. Sci Rep 9, 7324.

Wither, R. G., Lang, M., Zhang, L. and Eubanks, J. H. (2013). Regional MeCP2 expression levels in the female MeCP2-deficient mouse brain correlate with specific behavioral impairments. Exp Neurol 239, 49–59.

Xu, X., Garcia, J., Ewalt, R., Nason, S. and Pozzo-Miller, L. (2017). The BDNF val-66-met Polymorphism Affects Neuronal Morphology and Synaptic Transmission in Cultured Hippocampal Neurons from Rett Syndrome Mice. Front Cell Neurosci 11, 203.

Xu, X., Kozikowski, A. P. and Pozzo-Miller, L. (2014). A selective histone deacetylase-6 inhibitor improves BDNF trafficking in hippocampal neurons from Mecp2 knockout mice: implications for Rett syndrome. Front Cell Neurosci 8, 68.

Zhou, Z., Hong, E. J., Cohen, S., Zhao, W. N., Ho, H. Y., Schmidt, L., Chen, W. G., Lin, Y., Savner, E., Griffith, E. C. et al. (2006). Brain-specific phosphorylation of MeCP2 regulates activity-dependent Bdnf transcription, dendritic growth, and spine maturation. Neuron 52, 255–69.

